# Prediction of condition-specific regulatory maps in *Arabidopsis* using integrated genomic data

**DOI:** 10.1101/565119

**Authors:** Qi Song, Jiyoung Lee, Shamima Akter, Ruth Grene, Song Li

## Abstract

Recent advances in genomic technologies have generated large-scale protein-DNA interaction data and open chromatic regions for multiple plant species. To predict condition specific gene regulatory networks using these data, we developed the **Con**dition **S**pecific **Reg**ulatory network inference engine (**ConSReg**), which combines heterogeneous genomic data using sparse linear model followed by feature selection and stability selection to select key regulatory genes. Using *Arabidopsis* as a model system, we constructed maps of gene regulation under more than 50 experimental conditions including abiotic stresses, cell type-specific expression, and stress responses in individual cell types. Our results show that ConSReg accurately predicted gene expressions (average auROC of 0.84) across multiple testing datasets. We found that, (1) including open chromatin information from ATAC-seq data significantly improves the performance of ConSReg across all tested datasets; (2) choice of negative training samples and length of promoter regions are two key factors that affect model performance. We applied ConSReg to *Arabidopsis* single cell RNA-seq data of two root cell types (endodermis and cortex) and identified five regulators in two root cell types. Four out of the five regulators have additional experimental evidence to support their roles in regulating gene expression in *Arabidopsis* roots. By comparing regulatory maps in abiotic stress responses and cell type-specific experiments, we revealed that transcription factors that regulate tissue levels abiotic stresses tend to also regulate stress responses in individual cell types in plants.

## Introduction

Over the past decade, thousands of expression profiles have been generated using either RNA-seq or microarray hybridizations in the study of how environmental perturbations and developmental cues regulate gene expression in plants. Understanding the regulation of transcription in plants is crucial to improving crop productivity under challenging environmental conditions. However, computational tools in plant transcriptome analysis remain largely based on co-expression analysis (Serin et al. 2016), which may not be able to identify transcription factors (TFs) that control the expression of downstream genes for specific biological processes. Recent advancements in both *in vivo* and *in vitro* genomic experimental techniques have brought new data into the study of transcriptional regulation in plants. Large-scale ChIP-seq experiments (Song et al. 2016b), protein binding microarrays (Franco-Zorrilla et al. 2014) and DAP-seq experiments (O’Malley et al. 2016) have generated millions of candidate TF-target interactions. ATAC-seq and DNase hypersensitive assays have enabled profiling of activated chromatin regions under specific conditions and in specific tissue types (Lu et al. 2017; Maher et al. 2018; Cumbie et al. 2015; Zhang et al. 2012). A current major challenge is how to integrate protein-DNA interaction data and active chromatin region data with expression data to reveal fundamental regulatory mechanisms of gene expression under specific conditions in plants.

Conventionally, regulatory mechanisms were revealed by constructing genetic regulatory network (GRN) that are typically consisted of thousands of TF-target interactions. Many network inference approaches have been developed to construct GRNs by combining different types of genomic data. One notable example is mutual information (MI), which is a type of unsupervised machine learning approach that does not rely on any known interactions. Relevance Network (RN) is one of the first MI-based approach to infer interactions (Butte and Kohane 2000). To improve predictions, other MI-based inference methods were also developed (Margolin et al. 2006; Faith et al. 2007; Meyer et al. 2007). By contrast, supervised machine learning approaches which take known interactions as prior knowledge, have also been applied to the inference of network interactions. Several commonly used supervised models infer GRNs from expression data, including Support Vector Machine (SVM) (Mordelet and Vert 2008; Ni et al. 2016), Random Forest (RF) (Yip et al. 2012), Hidden Markov Model (HMM) (Ernst and Kellis 2010), least angle regression (LARS) (Haury et al. 2012), least absolute shrinkage and selection operator (LASSO) (Liu et al. 2014; Omranian et al. 2016), and elastic net (EN) (Altarawy et al. 2017). In spite of the success of these approaches, predictions solely based on expression profiles are far from perfect.

Other methods for inferring interactions focus on data types that present direct evidence of binding events. Binding site data has received much attention in recent years in the field of plant research as evidenced by databases such as PlantTFDB (Jin et al. 2016), AGRIS (Davuluri et al. 2003), and Grassius (Yilmaz et al. 2009) which have accumulated substantial amounts of data documenting experimentally identified binding sites. Previous studies have also identified a considerable number of binding sites from data obtained *in vivo* related to different environmental perturbations in plants. For example, binding sites were screened to construct regulatory networks in response to far red light (Chen et al. 2014b, 2014a), hormone (Fan et al. 2014; Song et al. 2016a; Shani et al. 2017) and fungal infection (Liu et al. 2015) in *Arabidopsis thaliana*. In contrast to expression data, binding site data present direct evidences of TF-target interactions. Based on available binding site data, several web-based tools have been developed to prioritize the targets of specific TFs for a group of genes using enrichment analysis. Some examples include TF2Network (Kulkarni et al. 2017) and Cistome (Austin et al. 2016), which compute enrichment of binding sites for corresponding TFs based on large collection of binding sites in *Arabidopsis*; PlantPAN 2.0 (Chow et al. 2016), which identified enriched combination of TFs for multiple plant species, and g:Profiler (Reimand et al. 2016), a tool designed to support binding site enrichment analysis of 213 species including 38 plant species.

However, direct experiments for binding site identification also have their limitations. For example, due to the high cost of ChIP-seq experiments, only a few TFs were typically screened under any specific condition. Compared to ChIP-seq, DAP-seq can identify possible targets of thousands of TFs efficiently (O’Malley et al. 2016). However, DAP-seq is an *in vitro* technique (Bartlett et al. 2017), and some binding sites detected by DAP-seq may not be available for binding under a given environmental perturbation. Therefore, integration of binding site and expression data is key to improving prediction accuracy under specific conditions or cell types.

In this study, we developed the **Con**dition **S**pecific **Reg**ulatory network inference engine (**ConsReg**), a machine learning approach which infers condition-specific regulatory networks from heterogeneous genomic data including expression data, DAP-seq data and ATAC-seq data. Training data were supplied to machine learning models to perform binary classification with feature selection by regularization. This procedure can prioritize and select the most relevant TFs for a specific environmental perturbation. We performed cross-validation for ConSReg using a compendium of expression data sets from 22 publications related to different environmental perturbations. The evaluation result shows that the features of the integrated representation can accurately predict expression of target genes (average AUC-ROC = 0.84).

Our analysis results highlight several important discoveries that provides new insights into the regulation of gene expression in plants. First, the appropriate selection of negative training datasets is crucial for the improvement of model performance, specifically, undetected genes are better negative training data than non-differentially expressed genes. Second, including ATAC-seq data significantly improved model performance regardless of the experimental conditions. Third, we found that the length of promoter regions contributes to the model performance. Although published studies show that stress regulated motifs are enriched in 500bp upstream of the TSS of target genes (Wang et al. 2012), our analysis showed that using 3KB upstream of TSS +0.5KB downstream of TSS as promoter provides better performance across all datasets (Kulkarni et al. 2017). When ConSReg was applied to data sets generated from drought, cold and heat perturbations, it successfully identified a pair of TFs, MYB44 and MYB77, that play active roles of co-regulating target genes in all three stresses. We also tested ConsReg in inferring cell type specific gene expression and cell type specific stress responses. We found that transcription factors that regulate stress responses in tissue level also regulate stress responses in individual cells. However, transcription factors that regulate cell type specific expression do not regulate stress responses.

Compared to other existing tools for regulatory network inference, one advantage of ConSReg is that the method does not require time course expression data and thus can be broadly applied in conditions where dynamic changes happens too early to collect time series data. This feature allows ConSReg to be used with emerging single cell sequencing data from plants to infer regulatory networks at single cell level. We tested ConSReg to *Arabidopsis* single cell RNA-seq (scRNA-seq) data of two root cell types (endodermis and cortex) and successfully identified five key regulators in two root cell types. Four out of the five regulators are supported by additional evidence from existing publications or cell type-specific expression data (See **Results**). Finally, we demonstrated that ConSReg has the potential to transform any published gene expression data into condition specific gene regulatory networks which will provide a system level overview of transcriptional regulation in plants. ConSReg is provided as a Python package and is available for download from GitHub (https://github.com/LiLabAtVT/ConSReg).

## Results

### Analysis overview

In this work, we focused on using protein-DNA interaction data and open chromatin data to predict the combinations of TFs that can best explain observed differential gene expression under different environmental perturbations or cell types. To achieve this goal, we have tested multiple machine learning methods in combination with different feature selection techniques to determine the optimal parameters and training strategies.

Our pipeline consists of two major steps (See **Figure 1A**). The first step is to integrate heterogeneous genomic data sets including interaction data generated from DAP-seq, open chromatin region data from ATAC-seq and expression data from RNA-seq/microarray experiments. This step also produced training, validation and testing data sets for machine learning models. The second step is to perform binary classification with sparse feature selection methods. The input feature matrix for classification was constructed from binding site information (DAP-seq) and activated chromatic regions (ATAC-seq) for a list of differentially expressed genes. These genes were obtained by standard statistical analysis approach using a contrast between a replicate group of treated samples and a replicate group of control samples. (See **Methods and supplementary tables** for more details).

**Figure 1.**
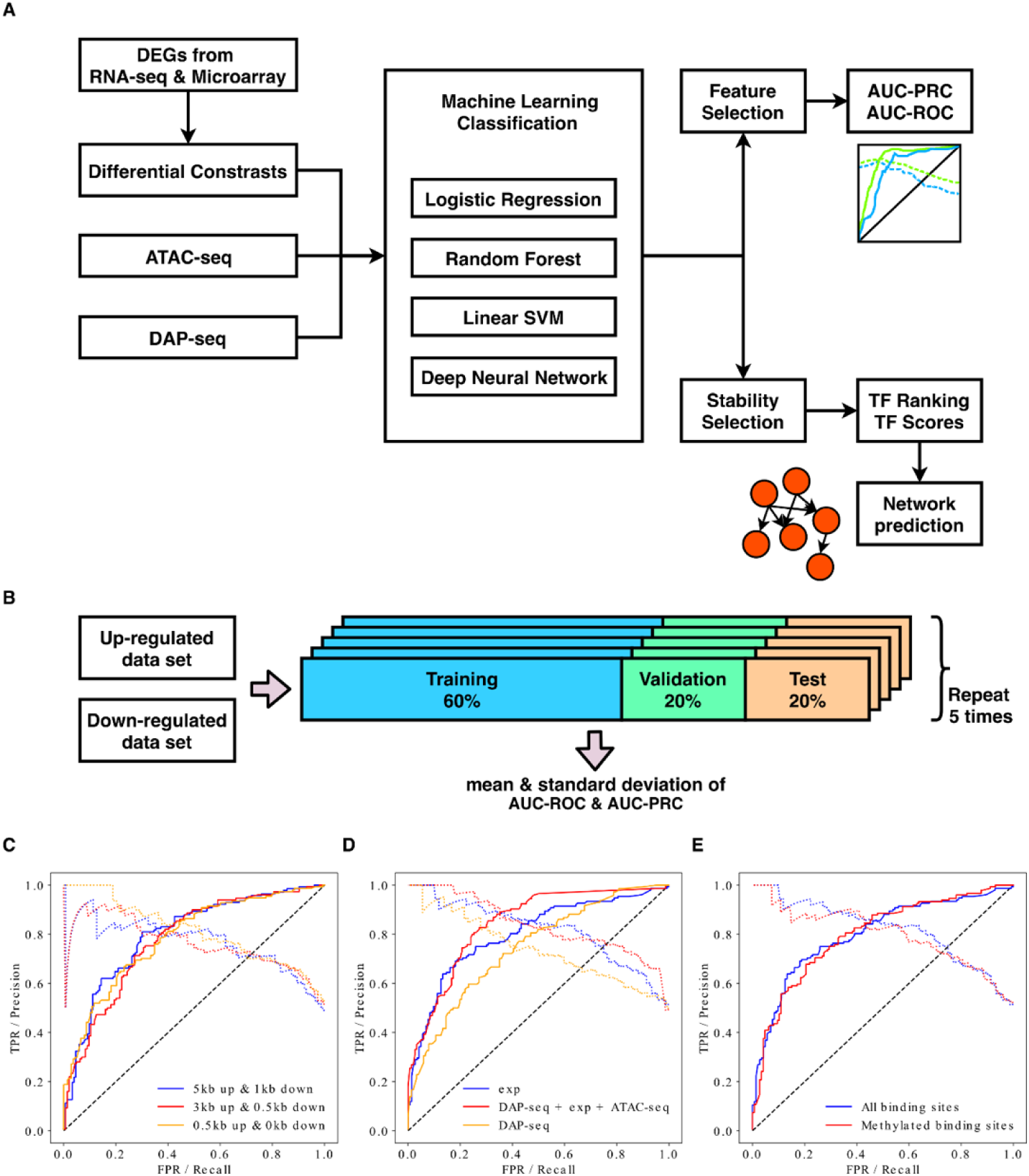
Prediction and evaluation. **A** The prediction and evaluation workflow. **B** Evaluation strategy. Cross-validation was performed. Each dataset was randomly split into three subsets, 60% for training, 20% for validation (hyperparameter tuning) and 20% for test. This was repeated five times for each data set. Mean and standard deviation of AUC-ROC and AUC-PRC were computed. **C, D, E** ROC and PRC curves for the performance of UR model trained using a differential contrast computed from a drought response dataset (differential contrast: PRJEB10930_3_T-PRJEB10930_3_C; see **supplementary table 1**) (Dubois et al. 2017). Different promoter region lengths were tested and compared in **C**. Different feature matrices were tested and compared in **D**. **exp**: feature matrices generated solely from expression data; **DAP-seq**: feature matrices generated solely from DAP-seq data; **DAP-seq + exp + ATAC-seq**: feature matrices generated from integrating all data types. Different types of DAP-seq binding sites were tested and compared in **E**. All curves were generated by averaging the false positive rates and false negative rates over five runs of cross-validation.

The goal of the machine learning method is to identify a minimum set of TFs that can best explain the observed gene expression data. Specially, we were using differentially expressed genes as positive training set to perform binary classification. For each gene, the feature matrix was consisted of all the interactions between TFs and their target genes. In our analysis, results from DAP-seq experiments were used to construct the feature matrix, however, other interaction data can also be included. Open chromatin regions were used to set a weight on the feature matrix. We analyzed up-and down-regulated genes separately, and we tested different methods of selecting negative training samples (see **Methods**).

**Table 1.** Number of interactions collected from each platform.

| Platform type | Number of interactions | Percentage |
| --- | --- | --- |
| DAP-seq | 1,812,475 | 97.11% |
| ChIP-seq | 50,513 | 2.71% |
| eY1H | 1,925 | 0.10% |
| Literature | 1,431 | 0.08% |
| Total | 1,866,344 | 100.00% |

Performances of machine learning models were evaluated by area under curve of the receiver operating characteristic curve (AUC-ROC) and area under curve of the precision-recall curve (AUC-PRC) computed from cross-validation. To prioritize important TFs for each condition, we assigned an importance score to each TF by performing stability selection (Meinshausen and Bühlmann 2010). We further explored the common combinatorial regulations in different environmental stresses (cold, heat, drought), by performing feature selection and co-regulators analysis.

We demonstrated the impact of different conditions on model performance, including using different negative training genes, selecting different promoter region length, integrating ATAC-seq data into the pipeline and using all or methylated DAP-seq binding sites. We generated ROC curves and PRC curves of the up-regulated (UR) feature matrix from one differential contrast as an example **(Figure 1 C D E**) to demonstrate the differences between these alternative setting for model training and feature construction. This example was selected from a recently published drought experiment (Dubois et al. 2017).

### Choice of negative training data sets is crucial to achieve better performance

As shown in a previous study, the choice of negative training data sets can significantly impact the performance of machine learning models (Natarajan et al. 2012). We systematically evaluated three different methods for selecting negative training genes and compared their performances for each model. These methods include: 1) non-significantly differentially expressed genes (**NDEGs**), which have p-value > 0.05; 2) low-expressed genes (**LEGs**), which have mean expression between 0 and 0.5; and 3) undetected genes (**UDGs**), which have a mean expression value equal to zero. The three methods were tested using evaluation dataset A (See **Methods**), where we constructed both an up-regulated (**UR**) feature matrix and a down-regulated (**DR**) feature matrix for each differential contrast. Machine learning models tested in this analysis include 1) logistic regression with lasso penalty (**LRLASSO**), 2) logistic regression with group lasso penalty (**LGLASSO**), 3) logistic regression with elastic net penalty (**LREN**), 4) logistic regression with Pearson Correlation Coefficient (**LRPCC**), 5) guided regularized random forest (**GRRF**), 6) linear support vector machine (**LSVM**).

For both UR and DR feature matrices, UDGs show consistently higher AUC-ROC than NDEGs and LEGs (**Supplementary figure 1A-C and 2A-C**). **Figure 2 A-C** shows one example of comparison between different negative training genes, using UR feature matrix (shaded shapes) and DR feature matrix (unshaded shapes) as input. The presented differential contrast was selected from a drought experiment (Dubois et al. 2017) (for evaluation results in more details, see **Supplementary Figure 3**). All these results show that, for both UR feature matrices and DR feature matrices, UDGs performed best among all three methods of selecting negative training genes. However, we do not find obvious difference for number of selected TFs among the three types of negative training genes (**Supplementary figure 1D-F** and **Supplementary figure 2D-F**). All models show very similar AUC-ROC value for both UR feature matrices and DR feature matrices.

**Figure 2.**
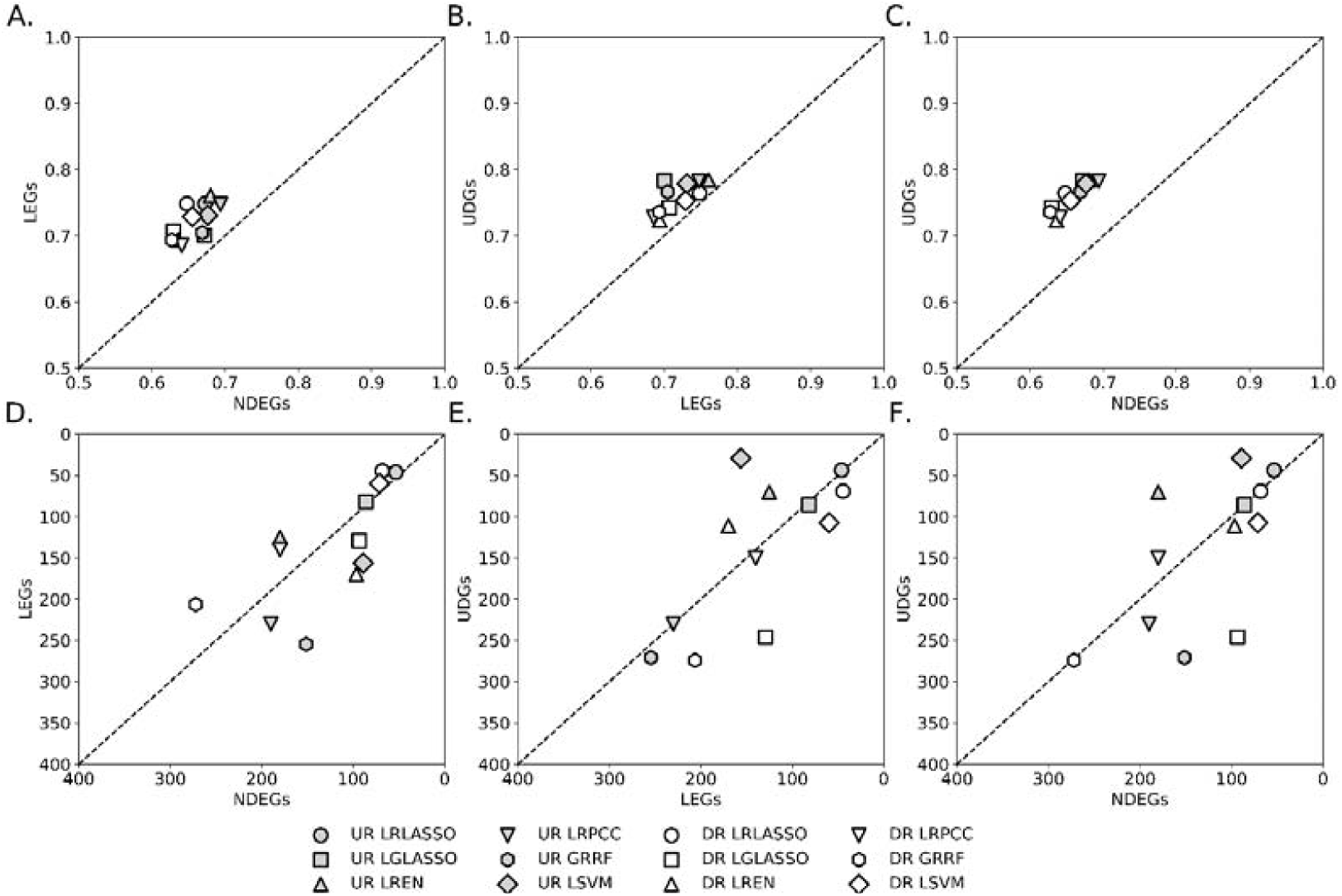
Performance evaluation for different classifiers and negative datasets. Evaluation results of one dataset (differential contrast: PRJEB10930_3_T-PRJEB10930_3_C; see **supplementary table 1**) was presented in the figure. This dataset was chosen from drought response experiment for *Arabidopsis* leaf tissue (Dubois et al. 2017). The full evaluation results from **evaluation dataset A** were shown in **supplementary figure 1** and **supplementary figure 2**. The following six classification strategies were tested. LRLASSO: logistic regression + LASSO; LGLASSO: logistic group LASSO; LREN: logistic regression + elastic net; LRPCC: logistic regression + Pearson correlation coefficient; GRRF: guided regularized random forest; LSVM: linear support vector machine. Three types of negative training genes were tested: 1) non-significantly differentially expressed genes (**NDEGs**); 2) low-expressed genes (**LEGs**); and 3) undetected genes (**UDGs**) (See **Methods** for more details). Each data point in the figure represents the average AUC value from the cross-validation (see **Methods**). **A, B, C** AUC values for different classifiers and negative datasets. **D, E, F** Number of selected TFs for different classifiers and negative datasets. Results generated from the UR feature matrices and DR feature matrices were marked by grey color and white color, respectively.

**Figure 3.**
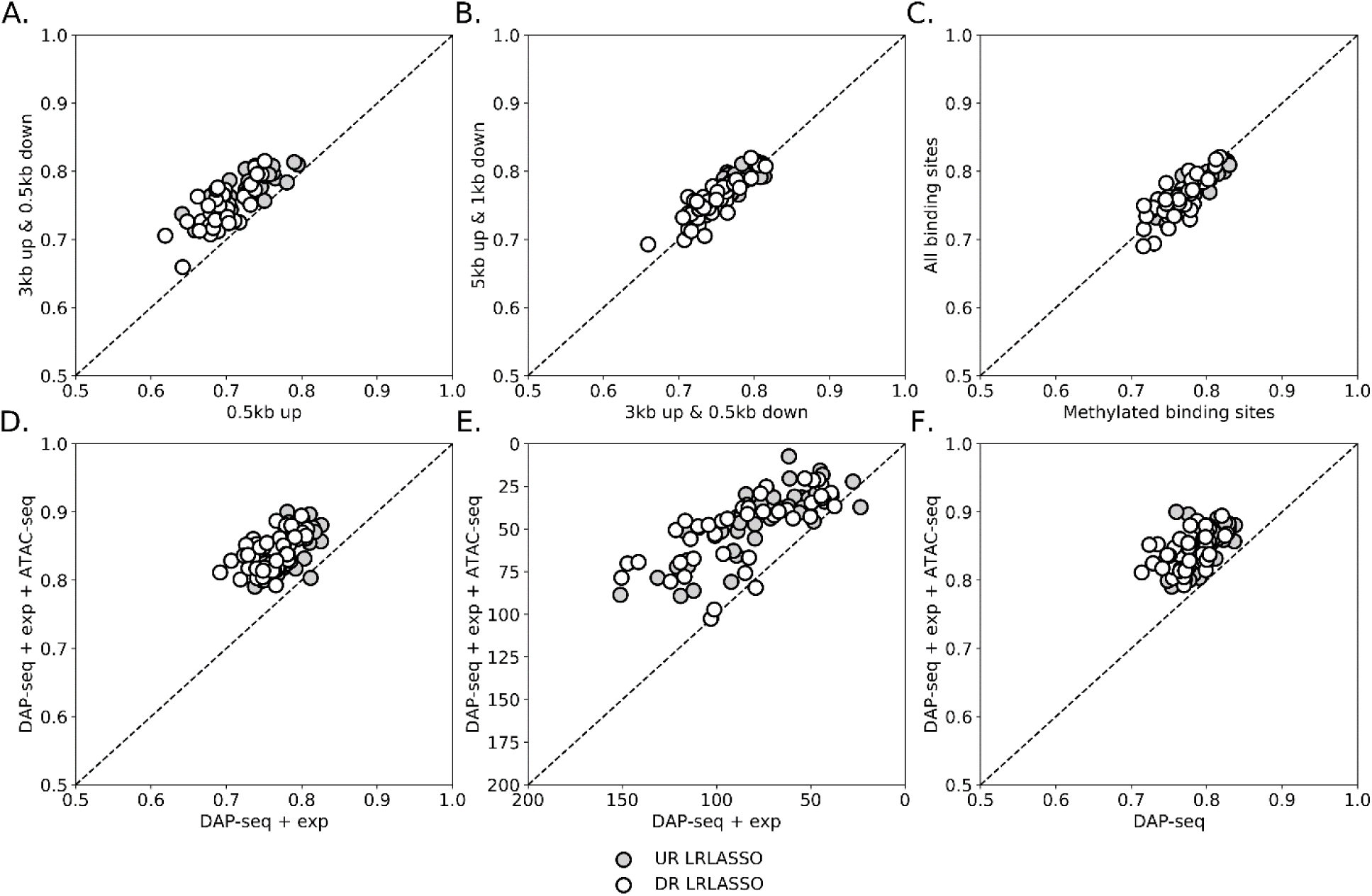
Performance evaluation for different feature construction methods. **A B,** AUC-ROC values of LRLASSO model when different promoter region lengths were tested using evaluation dataset B. **C**, AUC-ROC values for feature matrices constructed with all binding sites VS feature matrices constructed with methylated binding sites only. **D**, AUC-ROC values for feature matrices constructed with DAP-seq + expression (exp) + ATAC-seq data VS feature matrices constructed DAP-seq + expression. **E**, Number of selected TFs corresponding to the evaluation performed in **D**. **F**, AUC-ROC values for feature matrices constructed with DAP-seq + expression + ATAC-seq data VS feature matrices constructed with DAP-seq data only. The resulted feature matrices contain only numeric value zero and one (see **Methods**). For all subfigures, white circles represent evaluation result of UR model and shaded circle represent evaluation result of DR model.

Numbers of selected TFs from different machine learning models are quite different. LRLASSO selected fewer TFs than other methods, and standard deviation of the number of selected TFs is smaller than other models (**Supplementary figure 1J-L** and **Supplementary figure 2J-L**). Based on this observation, we performed following analyses using LRLASSO.

### Condition specificity of negative training genes

Although positive training genes in this study reflect condition-specific activities, it is unclear that whether negative training genes are also condition specific. One possibility is that all negative training genes are not detected under any condition tested. We checked whether UDGs are different in different environmental perturbations. For each differential contrast in each environmental perturbation, we computed the percentage of UDGs that are detected (fpkm > 0) in other perturbations. Then the percentages were averaged for each environmental perturbation. We found that this average percentage ranges from 72.54% to 91.76%, suggesting that UDGs in one condition are typically expressed in other environmental perturbation(s). Therefore, a large portion of UDGs remain inactive specific to one or multiple environmental perturbations (**Supplementary Figure 4**).

**Figure 4.**
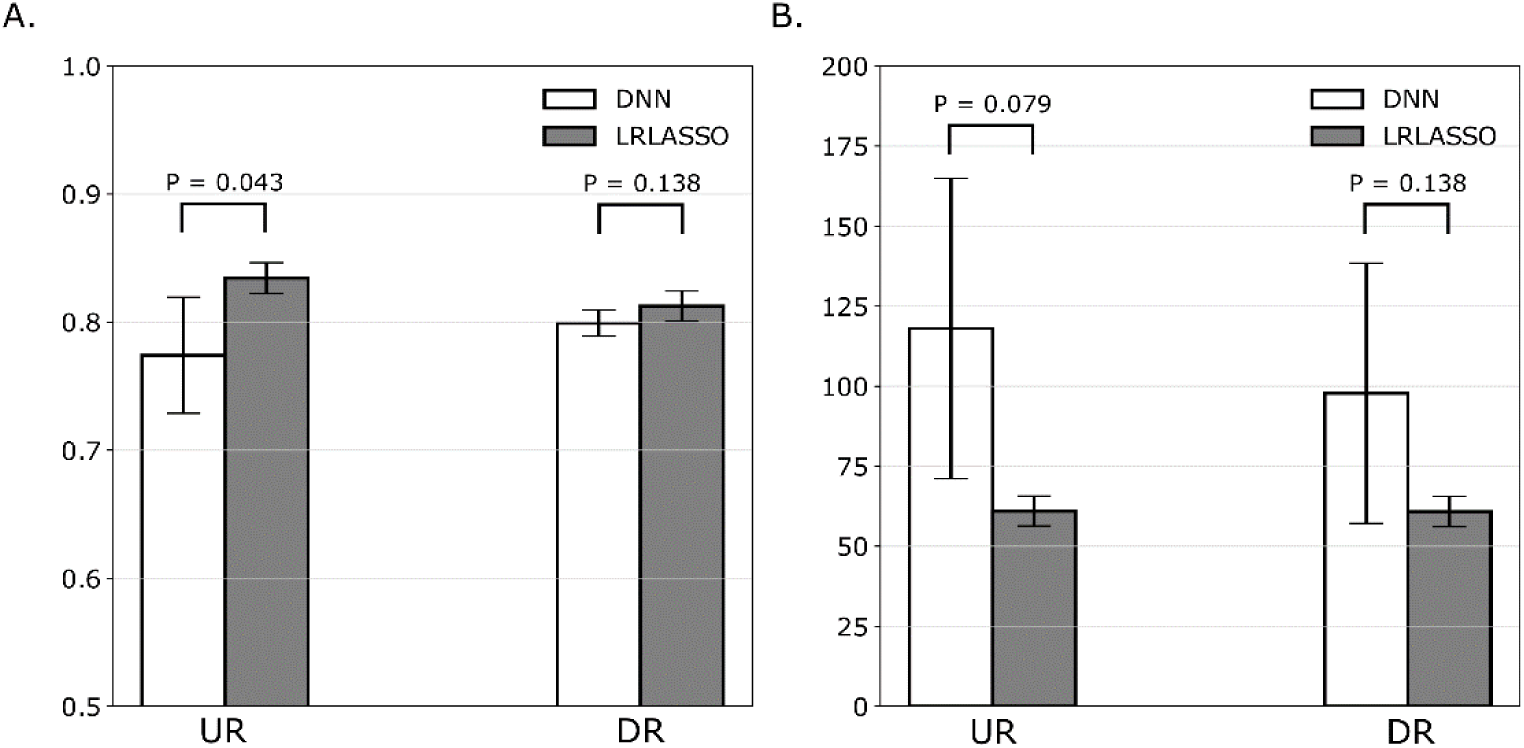
Comparison between LRLASSO and deep neural network (DNN). We compared the performance of LRLASSO and deep neural network (DNN). We used one differential contrast (GSE81202_18-GSE81202_16) to compute the AUC-ROC values of LRLASSO and DNN. **A**, Comparison of AUC-ROC values of LRLASSO and DNN. **B**, Comparison of number of selected TFs between LRLASSO and DNN. Error bars indicate standard deviations. P-values of significance were computed using Wilcoxon sum rank test and were marked above the bars.

### Choice of promoter region length affects model performance

TFs regulate expression of target genes by binding to regulatory elements located in the promoter regions of these genes. The combinatorial regulations of TFs have been studied in plants (Kaufmann et al. 2010). It has been shown that binding sites located within 5kb upstream region of transcription start sites (TSS) can better explain regulatory effect on the target genes than shorter regions (Kulkarni et al. 2017). However, the optimal promoter length has not been thoroughly investigated using condition-specific data. We therefore set the promoter region length up to 5kb upstream of TSS and 1kb downstream of TSS in feature construction step (See **Methods**) and tested different lengths under various environmental perturbations. We tested three types of promoter region: 1) 5kb upstream of TSS to 1kb downstream of TSS; 2) 3kb upstream of TSS to 0.5kb downstream of TSS; and 3) 0.5 kb upstream of TSS to TSS. **Figure 3 A**, **B** shows evaluation result of AUC-ROC values for three types of promoter regions evaluated on evaluation dataset B (See **Methods**). We observed consistent improvements when promoter region length was extended from 0.5 kb upstream to 3kb upstream + 0.5kb downstream. When the promoter region was further extended to 5kb upstream + 1kb downstream, no significant improvement was found. As shown in **Figure 3A** and **Figure 3B**, these results are consistent between both UR genes and DR genes (grey and white circles respectively). We plotted ROC curves and PRC curves of the UR genes from one differential contrast as an example **(Figure 1C**). The presented differential contrast was selected from a drought experiment (Dubois et al. 2017). In summary, our findings suggest that most of the binding sites predictive of gene expressions were successfully captured within 3kb upstream + 0.5kb downstream region.

### ATAC-seq data significantly improves the model performance

Although DAP-seq has shown higher throughput than earlier TF-target screening assay such as ChIP-seq (O’Malley et al. 2016), all DAP-seq binding sites are detected *in vitro*, and some of the binding sites in vitro might not be accessible in living cells. As suggested in a previous publication describing the DAP-seq assay, this limitation can be overcome by integrating DAP-seq with data on open chromatin regions (Bartlett et al. 2017). Therefore, we encoded open chromatin information from ATAC-seq data into the feature matrices (See **Methods**). To assess the impact of chromatin accessibility, the feature matrices were constructed either with or without integrating ATAC-seq data. We then compared the model performance of ATAC-seq included feature matrices to ATAC-seq free feature matrices using evaluation dataset B (See **Methods**). As shown in **Figure 3D**, for both UR and DR genes, there are consistent improvements when ATAC-seq data were included in the feature matrices. The other noticeable advantage of including ATAC-seq data is that it helped the model to select fewer TFs, producing a more interpretable result (**Figure 3E**). We further investigated whether including condition-specific expression and ATAC-seq data can better predict expression than using DAP-seq binding site information alone. To answer this question, we constructed feature matrices only by DAP-seq data (see **Methods**) and compared prediction results to feature matrices constructed with expression, ATAC-seq peaks and DAP-seq binding sites. The results show that including all three types of data has consistently improved performance (**Figure 3F**).

### Non-methylated binding sites do not improve the model performance

As described previously (O’Malley et al. 2016), DAP-seq can be performed in two ways: 1) sequence regular genomic DNA (gDNA), 2) sequence gDNA libraries in which methyl-cytosines were removed by PCR. The former is regular DAP-seq and the latter is called ‘ampDAP-seq’ (O’Malley et al. 2016). We tested the performance of two sets of binding sites: 1) using all available DAP-seq binding sites, 2) using only methylated DAP-seq binding sites. Although it was reported that many DAP-seq binding sites (∼180,000) were occluded by DNA methylation, our result shows that, compared to using all available DAP-seq binding sites, using ampDAP-seq binding sites does not provide better prediction result (**Figure 3C**).

### Comparison between LRLASSO and DNN

In recent years, deep neural network (DNN) has been extensively applied in the field of genomics to model gene regulations (Alipanahi et al. 2015; Zhou and Troyanskaya 2015; Singh et al. 2016). We further explored whether DNN can bring better performance than LRLASSO. A previous study has introduced a DNN-based feature selection method (Li et al. 2015). We used a similar strategy in our analysis (See **Methods**) to prioritize TFs and compare the result to LRLASSO. Since DNN usually needs large number of training examples to estimate thousands of parameters, we selected a differential contrast which has the largest number of training examples (8,948 genes for UR and DR feature matrices, respectively). Shown in **Figure 4 are** AUC-ROC values and selected number of features from five rounds of cross-validation. For UR feature matrix, the performance of LRLASSO is significantly better than DNN (**Figure 4A**, Wilcoxon rank-sum test, p-value = 0.043), whereas the performances do not show significant difference for DR feature matrix (**Figure 4A**, Wilcoxon rank-sum test, p-value = 0.138). LRLASSO selected fewer TFs than DNN (**Figure 4B**) and has more robust performance compared to high variation of number of selected TFs from DNN. Given these findings, we concluded that DNN-based feature selection does not perform better than LRLASSO in our analysis.

### ConSReg outperforms simple enrichment test

An enrichment test has been applied in recent studies to prioritize TFs given a set of input genes (Kulkarni et al. 2017; Reimand et al. 2016; Chow et al. 2016; Austin et al. 2016). However, enrichment-based prediction does not take into consideration combinations of multiple TFs. We compared our prediction pipeline to an enrichment-test-based method (See **Methods** for details). For both ConsReg and enrichment test method, we computed AUC-ROC values using evaluation dataset B (See **Methods** for details). As shown in **Figure 5**, ConsReg outperforms enrichment test in all tested differential contrasts. AUC-ROC values for ConsReg are significantly higher than enrichment test (Wilcoxon rank-sum test, p-value = 7.61 × 10^−9^ for UR feature matrices and p-value = 7.60 × 10^−9^ for DR feature matrices).

**Figure 5.**
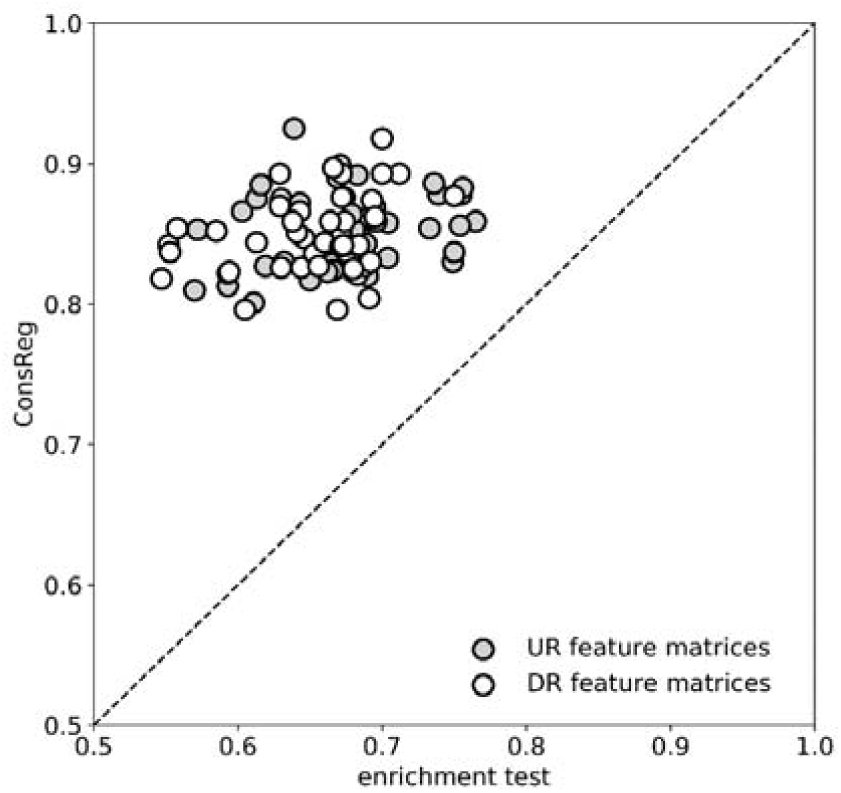
Comparison between ConSReg and enrichment test. We used LRLASSO as machine engine of ConSReg to predict gene expressions. We compared the performance of ConSReg to an enrichment-based method (see **Results** and **Methods**) using evaluation dataset B. AUC-ROC values for both methods were plotted in this figure (see **Methods** for details). White circles represent evaluation result of UR model and shaded circle represent evaluation result of DR model.

### ConSReg successfully identified known TFs that regulate abiotic stress

We performed comprehensive investigation of active TFs in multiple abiotic environmental perturbations using our prediction pipeline. ConSReg was ran for all differential contrasts included in evaluation dataset B, which encompasses nine common environmental perturbations: cold, heat, drought, salt, wounding, osmotic stress, red light, blue light and high light. For each differential contrast in each environmental condition, we assigned importance score to each TF by LRLASSO + stability selection (See **Methods**). The highest importance score was selected as representative importance score for each TF in each environmental condition. We then inferred a GRN for each environmental condition (See **Methods**). We computed the basic network properties of these GRNs (**Supplementary table 2**). We counted how many times each TF achieved a score higher than 0.5 across nine environmental conditions. This number is hereafter referred as ‘condition count’ which was then used to rank all TFs.

MYB and ERF protein families are known for regulating many abiotic stress responsive genes (Fujita et al. 2006; Müller and Munné-Bosch 2015). We found presence of many ERF and MYB/MYB related TFs in our top 20 candidates generated from UR feature matrices, including five TFs from MYB/MYB related family (AT1G18330, AT3G50060, AT1G49010, AT5G67300 and AT1G74650) and two TFs from ERF family (AT2G31230 and AT4G16750).

Some of the known stress related TFs were ranked as top candidates by condition count (See **Supplementary Table 3**). For example, AT1G27730 (ZAT10), which was reported to be involved in defense response of plants to abiotic stresses such as heat, cold, drought and salt (Sakamoto 2004; Mittler et al. 2006; Xie et al. 2012), was assigned high condition count in both UR and DR feature matrices (condition count = 8 for both UR and DR feature matrices). Our result is consistent with a previous study which suggests that ZAT10 may function as both a positive and a negative regulator of defense response to abiotic stresses (Mittler et al. 2006). More interestingly, our result indicates that ZAT10 is likely to be an active regulator during multiple light environment perturbations including high light, blue light and red light. However, the involvement of ZAT10 in multiple light stresses has not yet been well characterized. Although there are evidences suggesting that ZAT10 mediates a response to high light (Rossel et al. 2007; Gordon et al. 2012), little evidence has been reported about the involvement of ZAT10 in red light and blue light response. A previous study identified ZAT10 as the substrate of Mitogen-Activated Protein Kinase (MAPK) and showed that ZAT10 can directly interact with two MAPKs: MPK3 and MPK6 (Nguyen et al. 2012). It has been reported that MAPKs can be activated by blue light to modulate the response (Johnson and Lapadat 2002). Given the evidences above, we hypothesize that ZAT10 may affect the blue light response by interacting with blue-light-induced MAPKs.

The other notable example of a gene with high condition count (condition count = 8) is AT2G46680 (ATHB7), a member of the homeodomain-leucine zipper family (HD-ZIP). ATHB7 was already shown to confer salt and drought tolerance to plants (Olsson et al. 2004; Mishra et al. 2012; Pruthvi et al. 2014) and act as a negative regulator for plant growth (Olsson et al. 2004). In contrast, its functional role for other stress responses is less well characterized. Our result shows that ATHB7 is an active regulator for both UR genes and DR genes, suggesting it may function as both positive and negative regulator for multiple environmental stresses (See **supplementary table 3**).

### ConSReg uncovers combinatorial regulations

TFs are known to modulate expression of target genes by combinatorial regulation in plants (Singh 1998; Kaufmann et al. 2010). Combinatorial regulations between TFs can be established either by forming protein complexes between TFs, or indirect interactions between TFs (Kaufmann et al. 2010). We explored possible combinatorial regulations between TFs with high importance score (> 0.5) for three environmental perturbations related to abiotic stresses (cold, heat, drought). For each environmental perturbation, we computed importance scores and inferred condition-specific GRN (See **Network Inference** in **Methods**). Due to the large size of each GRN, we selected the TFs from each GRN and visualized only the subnetworks formed by these TFs. Shown in **Supplementary figure 5** and **Supplementary figure 6** are subnetworks of TFs for cold, heat and drought condition. To better understand this hierarchy of regulation, we clustered the TFs using a simulation-annealing-based algorithm (Gerstein et al. 2012). We enforced the algorithm to cluster TFs in three clusters: 1) top TF cluster, in which TFs have many out-going edges and less in-coming edges from other TFs, 2) bottom TF cluster, in which TFs have many in-coming edges and less out-going edges from other TFs. 3) intermediate TF cluster, in which TFs have balanced out-going edges and in-coming edges from other TFs. The visualization result shows that many TFs were put into top TF cluster, suggesting that many of these TFs serve as master regulators regulating fewer number of intermediate TFs and bottom TFs (**Supplementary figure 5** and **Supplementary Figure 6**).

**Figure 6.**
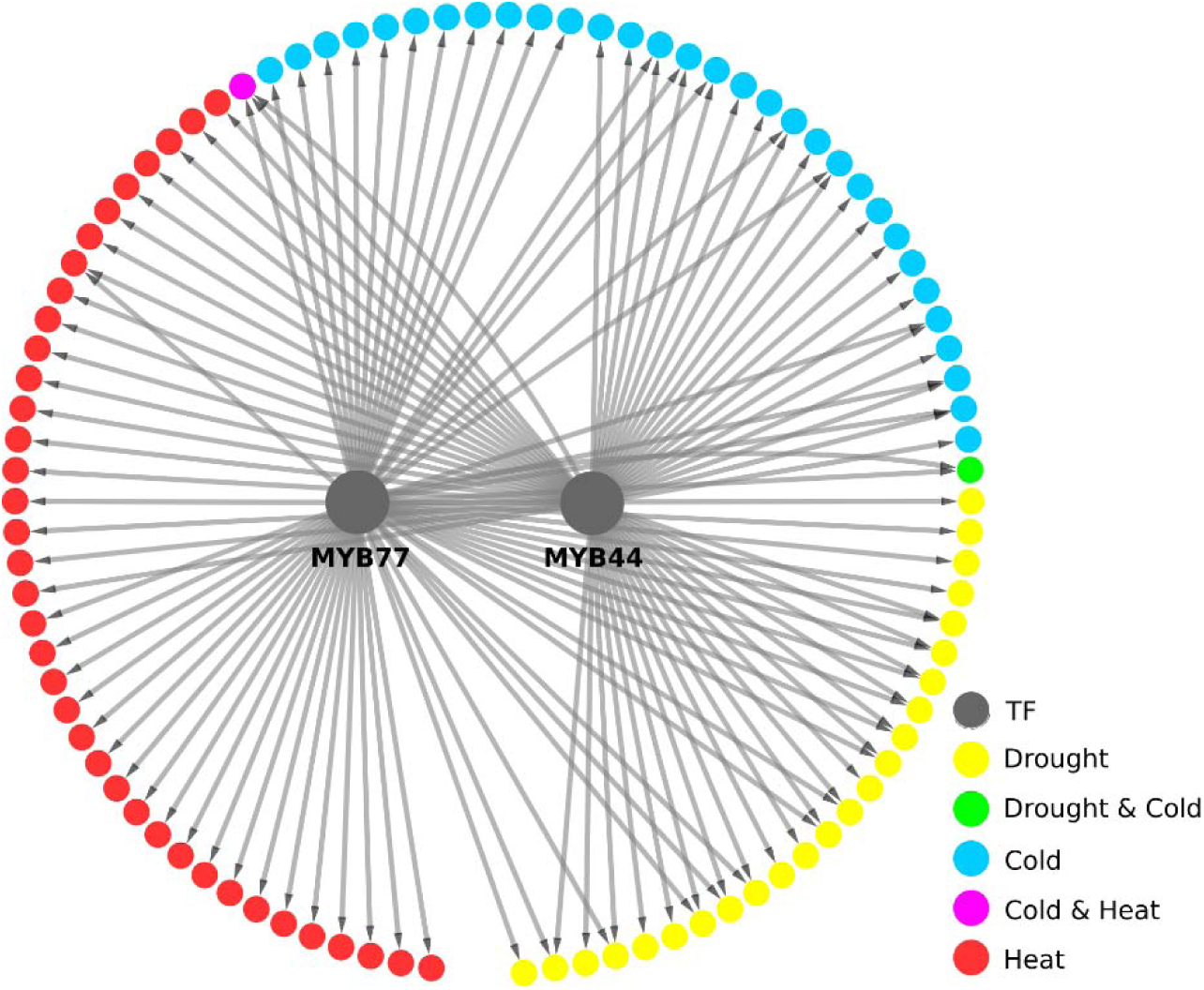
Combinatorial regulation between MYB77 and MYB44 and their top target genes. Located in the center are two regulators MYB77 and MYB44. Their respective top 20 DEG targets were plotted along the circle. DEGs in each environmental condition were sorted by ascending order of p-value. DEGs were marked by different color(s) to indicate their associated environmental condition(s).

We identified co-regulating modules of TFs from each GRN using a previously published tool CoReg (Song et al. 2017). Then we identified maximum common co-regulating TFs (See **Methods**) across the three conditions. While no common co-regulating TFs can be found for UR GRNs, we found a pair of common co-regulating TFs for DR GRNs: MYB77 (AT3G50060) and MYB44 (AT5G67300). We plotted the network of MYB77 and MYB44 with their respective top 20 DEG targets in each condition sorted by ascending order of p-value. (**Figure 7**). It was previously reported that MYB77 and MYB44 may function redundantly to enhance auxin signaling (Zhao et al. 2014). In plants, auxin signaling is involved in stress responses by regulating plant growth (Park 2007). Although involvement of MYB77 and MYB44 in stress response has not been well characterized, our result suggests they may modulate stress responses through combinatorial regulation.

**Figure 7.**
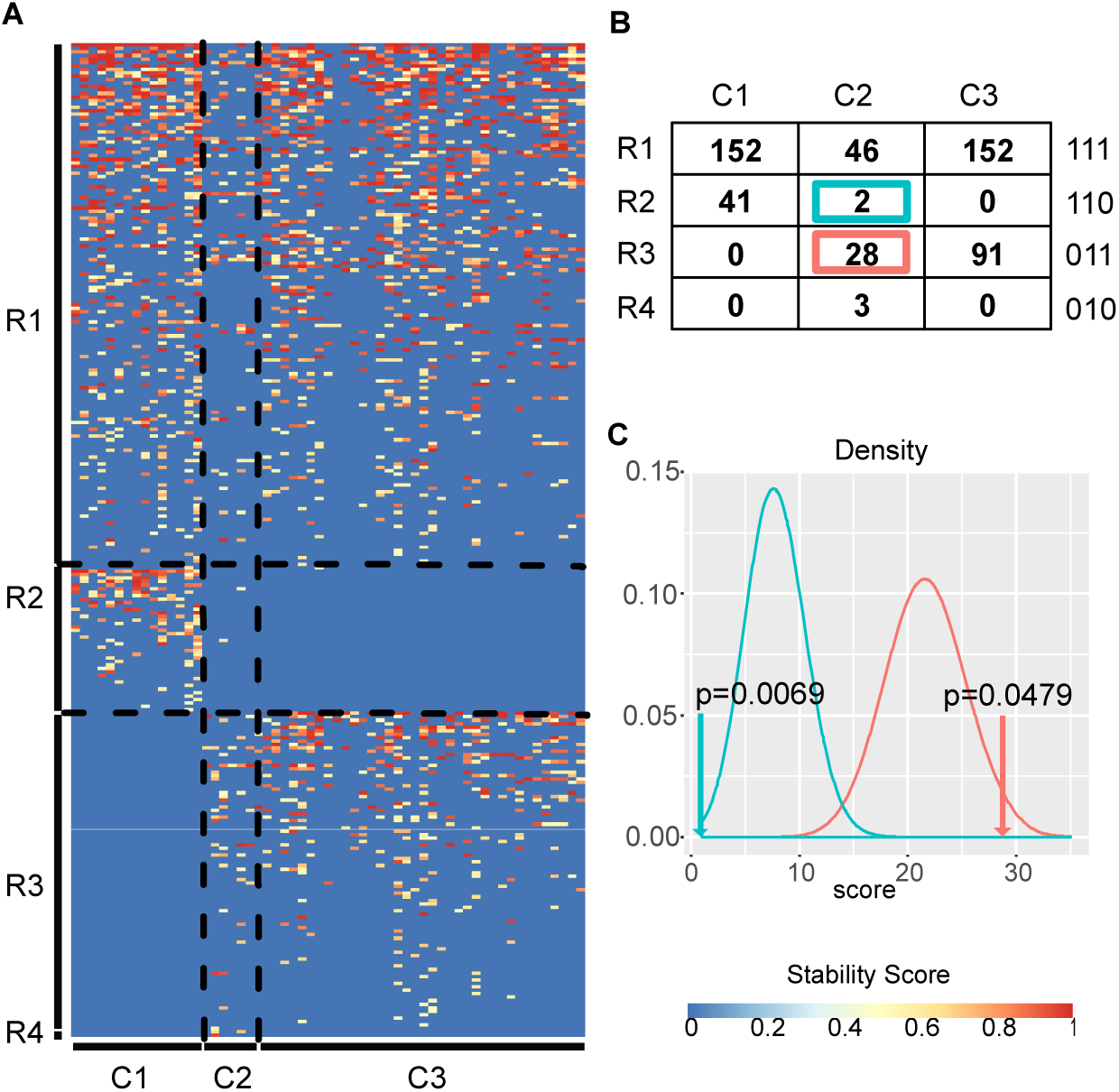
(A) Heatmap of stability scores. Each row represents one TF and each column represents one experimental condition. R1. TFs regulate both cell type specific expression and stress responses. R2. TFs mostly regulate cell type specific expressoin. R3. TFs do not regulate cell type specific responses. R4. TFs only regulate cell type specific responses. C1. Cell type specific experiments. C2. Cell type specific salt stresses experiments. C3. Abiotic stress experiments in tissue or organ level. (B) Number of genes belong to each category. (C) Simulated density distribution of number of genes in the R2C2 category (green) and R3C2 category (red). Arrow heads indicate the observed numbers. P values are calculated based on 100,000 random sampling.

### ConSReg reveals hidden rules of combinatorial regulation in plant stress response

In multi-cellular organisms, one central problem is to understand how cell type-specific responses were achieved through the combintorial regulation of transcription factors. Earlier results using cell types specific salt-induced gene expression in Arabidopsis roots have suggested that there are unknown, cell type-specific stress regulatory genes that mediate cell type specific responses (Dinneny et al. 2008). However, very limited genome wide binding site were available when the gene expression data were generated. To test this model, we applied ConSReg to these published gene expression data and predicted regulatory networks. We downloaded cell-type-specific expression data for *Arabidopsis* root (Li et al. 2016) (See **Methods**) and *Arabidopsis* root cells under salt stress (Dinneny et al. 2008). ConsReg integrated the two expression data sets with DAP-seq data and ATAC-seq data as described above (See **Methods**). An importance score for each TF was computed and GRN for each cell type was inferred. Compared to the GRNs of environmental perturbations, these GRNs have lower average number of nodes, edges and active TFs (See **supplementary table 2**).

The stability scores are visualized in a heatmap (Figure 7A). We found these TFs can be classified into four categories. A TF is predicted to regulate gene expression under one experimental condition if the stability score of this TF is higher than 0.5. We found that 152 TFs (Figure 7A, group R1) regulate both cell type-specific gene expression and abiotic stress responses. Among these 152 TFs, 46 are also found to regulate cell type-specific stress responses. We found 41 genes (group R2) only regulate cell type-specific gene expression, but not stress responses, whereas 91 genes (group R3) only regulate stress responses but not cell type-specific gene expression. Surprisingly, amont the 41 genes that regulate cell type-specific gene expression, only two TFs are also regulate cell type specific stress response. This number is statistically significantly smaller than expected by random chance (p = 0.0069). In contrast, among 91 TFs that regulate stress responses in bulk tissues, 28 TFs are also regulate cell type-specific stress responses, which is statistically significantly more than expected by random chance (p = 0.0479). This result shows that TFs play roles in regulate stress responses are also more likely to be involved in regulate cell type-specific stress responses. On the contrary, TFs that regulate cell type-specific expression tend to not regulate stress responses in individual cell types.

### Inferred regulatory genes from single cell RNA-seq data agree with bulk sequencing results

We applied ConSReg to *Arabidopsis* scRNA-seq data of two root cell types (endodermis and cortex). scRNA-seq data was generated by drop-seq assay in a recent study (Shulse et al. 2018). UR and DR feature matrices were generated by comparing cortex cell type to endodermis cell type and importance scores were computed using these feature matrices (See **Methods** for details). TFs with importance score > 0.5 were considered as predicted regulator. Among the predicted regulators of single cell gene expression from cortex cluster and endodermis cluster, we found 4 and 5 genes are consistently predicted (stability score > 0.5) as regulators for genes preferentially expressed in cortex and endodermis clusters respectively. Among these genes, we found two genes (ATWRKY27 and ATHB34) are commonly found in both cortex and endodermis clusters. The two commonly predicted factors (AtWRKY27 and ATHB34) could represent common regulators for both cortex and endodermis cells. Cortex cluster contains a unique regulator gene, ERF115, whereas endodermis cluster contains BBX31 and TGA6. Surprisingly, among more than 20 ERF genes included in our input dataset, our model selected ERF115, which is known to regulate cell cycle in QC cells (Heyman et al. 2013). Although QC cells are not cortex or endodermis cells, this discrepancy could due to the noise in assigning single cell sequencing data to specific cell types. Some QC cells may have similar expression profile as young cortex or endodermis cells. For example, the commonly used SCR marker include cells from both QC and endodermis. The BBX31 gene is not only predicted as regulator for single cell gene expression data, but is also found to be regulator in bulk RNA-seq (importance score = 0.98 in E30 cell type, see **supplementary table 4**), strongly supporting the role of this gene in controlling gene expression in endodermis. The other endodermis predicted regulator, TGA6, is known as a regulator that mediate Phytoprostanes inhibition of root growth (Mueller et al. 2008). The ATHB34 gene is known to be highly expressed only in stele and pericycle, the predicted role of ATHB34 suggest that this gene could be another candidate gene that move from one cell type to another to regulate fate of neighboring cell types. Finally, AtWRKY27 is predicted as a regulator only in developing cortex (importance score = 1), and have minor role in developing endodermis or maturing endodermis (importance score ∼0.2). This result suggests the cortex population from single cell experiment could be more similar to developing cortex cell in bulk RNAseq results, whereas the endodermis population in single cell experiment is a mix of both developing and maturing endodermis cells.

## Discussion

### Including other genomic features in ConSReg

Apart from the genomic features we tested in this work, our prediction pipeline can be easily extended and applied to other types of expression data. The limitation of DAP-seq data is that some of the interactions detected by DAP-seq may not be active under specific *in vivo* conditions (Bartlett et al. 2017). One solution to address this issue is to integrate DAP-seq data with more genomic features that confer *in vivo* binding specificity of TFs. Such genomic features may encompass 1) the activities of TFs and their target genes and 2) the chromatin accessibility for TF binding sites. The first criterion can be satisfied by including expression of TFs and targets in feature matrices. There are abundant expression data sets generated for model plant species *Arabidopsis* under different environmental perturbations, including drought (Dubois et al. 2017), heat (Rawat et al. 2015; Kilian et al. 2007), cold (Schlaen et al. 2015; Gehan et al. 2015; Kilian et al. 2007), and salt stresses (Suzuki et al. 2016; Kilian et al. 2007). For the second criterion, ATAC-seq is used in the current analysis (Lu et al. 2017). However, this ATAC-seq data is generated for seedling and roots. If additional tissue or condition specific data become available, our method can integrate these new genomic features. Other experimental approaches such DNase-seq and MNase-seq (Bartlett et al. 2017) can also be used in the place of ATAC-seq experiments.

### Apply to single cell expression data

The recent advancement of sequencing technology has made it possible to investigate gene expressions at the single cell level. We have demonstrated application to scRNA-seq data in our analysis. To improve our current method, an important issue to be addressed is the sparsity of single cell data, which is usually characterized by zero-inflated read counts for majority of genes (Vallejos et al. 2017). A large portion of zero read counts may arise from technical noise or biological variability between single cells (Vallejos et al. 2017). These phenomena are known as ‘dropout’ events. In previous work, attempts were made to address stochastic dropout by modeling it as a three component mixture model (Kharchenko et al. 2014), two-component mixture linear model (Finak et al. 2015) or exponential function of expected expression (Pierson and Yau 2015).Our current method uses variable genes generated by Seurat (Butler et al. 2018) as positive genes, which were selected by finding the outliers on a mean variability plot. However, this selection process is based on normalized expressions which are not generated with error model that explains dropout events. In future work, error model discussed in previous studies can be included in ConSReg to obtain better positive training genes.

### Potential future improvement

While ConsReg achieved good performance (average ROC-AUC = 0.84), the tool can be further improved by either enhancing the model performance or by adding more functionalities that provides more detail concerning dynamic regulation. Open chromatin regions have been reported to be both cell-type-specific (Boyle et al. 2008; Thurman et al. 2012) and condition-specific (Hesselberth et al. 2009) as revealed by distribution of DNaseI hypersensitive sites (DHSs). In our analysis, expression data and ATAC-seq data were not generated under the same conditions nor from the same tissue type. This is because only data from roots and seedlings are currently available for Arabidopsis (Lu et al. 2017). We merged all open chromatin regions detected in two tissue types to maximize the discovery of potential interactions. This could introduce false positives and compromise the ability of the model to predict condition-specific interactions. In the future, such false positives can be reduced by integrating open chromatin data and expression data generated under the same conditions and same tissue type, as more data accumulate.

Another improvement could be made by altering the assumption of LRLASSO. Currently, our LRLASSO model assumes that the combinatorial regulations among TFs are identical for all DEGs (trained coefficients are identical for all DEGs). However, compared to real regulations, this is a simplistic assumption. To address this issue, we can either use information from other data types or cluster genes and adaptively fit local linear model to each cluster. In this case, the inferred combinatorial regulations may better represent the truth. We will leave this to future exploration in our follow-up work.

### Additional possible functionalities

To better understand how condition- / cell-type-specific regulation changes across different condition/cell type, networks inferred by ConsReg can be compared. For example, when applied to single cell expression data, or bulk expression data with many time points, network comparisons can identify different regulation at different time points and how a given network dynamically changes over a time series. This would allow us to capture transient and dynamic combinatorial regulations. For cell-type-specific expression data, an effective strategy might be to investigate the specificity of network module(s) for each cell type or a group of cell types. Modules found in many cell types may characterize fundamental pathways and modules highly specific to few cell types may play unique functional roles.

## Conclusions

In this study, we developed a novel computational tool, ConSReg. We performed comprehensive analyses to identify the factors that affect the performance of machine learning models and the optimal settings for constructing feature matrix. We showed that ConSReg outperforms current enrichment-test-based tools in the task of predicting gene expression. Network analysis for the GRNs inferred by ConSReg revealed a novel combinatorial regulation between MYB44 and MYB77 in response to cold, heat and drought stresses. We applied ConSReg to *Arabidopsis* scRNA-seq data of two root cell types (endoderims and cortext) and successfully regulators supported by existing publications. Finally, we showed that TFs regulate stress responses at whole tissue levels tend to also regulate stress responses at individual cell types. ConSReg provides a useful way of integrating currently available genomic features with published gene expression data to infer regulatory networks and to better understand mechanisms of gene regulation.

## Methods

### Bulk RNA-seq and microarray expression data

We re-analyzed published RNA-seq and microarray expression data for *Arabidopsis* from 20 experiments which were generated under different environmental and hormonal perturbations including cold, heat, drought, wounding, salt, osmosis, blue light, high light, far red light, abscisic acid (ABA), salicylic acid (SA), jasmonic acid (JA), auxin and thermospermine oxidase (T-Spm) (See **supplementary table1**). We also re-analyzed cell-type-specific gene expression in *Arabidopsis* root generated in our previous publication (Li et al. 2016), and gene expression of cell-type-specific responses to salt stress in *Arabidopsis* root (Dinneny et al. 2008) (See **supplementary table1**). The latter data set was generated using microarrays. We downloaded the pre-analyzed data files from the EBI expression atlas (Kolesnikov et al. 2015). Processed data is available in the ArrayExpress database (http://www.ebi.ac.uk/arrayexpress) under accession number “E-GEOD-7641”. We compiled **differential contrasts** for each experiment. Each differential contrast was defined as the contrast between a replicate group of treated samples and a replicate group of control samples (**See supplementary table1**). We used whole root samples as the control group for the cell-type-specific expression experiment because no environmental perturbations were used in the experiment. For RNA-seq data, we used a published protocol to identify differentially expressed genes. In brief, we used STAR for read mapping, featureCounts for read counting and DESeq2 for differential expression analysis. Differentially expressed genes (DEGs) were identified as genes with FDR < 0.05. Microarray data were analyzed using protocol established at the EBI expression atlas. For both RNA-seq and microarray data, we used the average gene expression level across all the samples within one replicate group.

### scRNA-seq expression data

We re-analyzed published *Arabidopsis* scRNA-seq data for two root cell types (endodermis and cortext) (Shulse et al. 2018). Expression data was downloaded from Gene Expression Omnibus (GEO) with accession number GSE116614 (https://www.ncbi.nlm.nih.gov/geo/query/acc.cgi?acc=GSE116614). The downloaded expression matrix contains read counts of each gene for each cell. Expression matrix was filtered by selecting cells that have more than one expressed gene and genes that are expressed in more than one cell. Read counts were normalized using log normalization method in the Seurat package (Butler et al. 2018). The normalized expressions were used to cluster cells by a graph-based clustering approach in the Seurat package. The identified clusters were assigned with known *Arabidopsis* root cell types by computing index of cell identity (ICI) scores between each cluster and the profile of marker genes generated by a previous study (Efroni et al. 2015). ICI score characterizes the probability that each cluster represents a known root cell type. We assign the cell type of highest probability to the cluster. Next, we used clusters identified as endodermis and cortex cells to compute fold change of each gene. SCDE package (Kharchenko et al. 2014) was used to fit an error model of dropout events and compute fold change of each gene with respect to the comparison of cortex VS endodermis.

### TF-target interaction and open chromatin data

We downloaded BED files of peak regions for 387 TFs from a published *Arabidopsis* DAP-seq dataset (O’Malley et al. 2016). All BED files are available at Plant Cistrome Database (http://neomorph.salk.edu/dap_web/pages/index.php). Interactions for the DAP-seq dataset were generated by assigning corresponding genes to each peak using R package ChIPseeker (Yu et al. 2015). When the promoter region was set as 5kb upstream and 1kb downstream region of TSS, a total of 1,812,475 interactions were identified from DAP-seq peaks. Among these interactions, 1,540,984 interactions were generated from regular DAP-seq, where methylations were not removed from genomic DNA and 1,280,138 interactions were generated from amp-DAP-seq, where methylation were removed. These two types were merged, which resulted in a total of 1,812,475 non-redundant interactions. We also compiled a list of published TF-target interactions generated by literature curation and ChIP-seq (Jin et al. 2016), eY1H (Taylor-Teeples et al. 2014; Sparks et al. 2016; Shani et al. 2017), and other methods. All interactions were merged and duplicates were removed, which resulted in a set of 1,866,371 interactions. The total number of interactions provided by DAP-seq account for 97.11% of the interactions (See **table1**), therefore we only used the DAP-seq data for simplicity of data integration and feature construction. For the open chromatin region data, we downloaded the *Arabidopsis* ATAC-seq peaks from a published study which identified open chromatin regions for whole seedlings and roots (Lu et al. 2017). All BED file for the peaks are downloaded from GEO website (https://www.ncbi.nlm.nih.gov/geo/query/acc.cgi?acc=GSE85203).

### Evaluation dataset A and evaluation dataset B

we constructed different evaluation datasets. The reason for using different evaluation datasets is to provide sufficient positive/negative training genes for machine learning and feature selection methods. For all expression experiments, we selected differential contrasts which provide more than 500 positive and 500 negative genes for all three types of negative genes (NDEGs, LEGs, UDGs). We compiled these differential contrasts into evaluation dataset A (**supplementary table 1**). After we determined that UDGs are better negative training sets than other two tpyes of training sets, we selected differential contrasts which provide more than 500 positive and 500 negative genes for only UDGs and compiled them into evaluation dataset B. This data set was then used to evaluate the performance of integrating ATAC-seq data and performance of different types of DAP-seq data.

## Feature construction

Based on expression data, we constructed differential contrasts between replicate group of control and treatment samples. Each replicate group typically include expression data of multiple samples and each differential contrast produced a list of genes with fold change, mean expression value and p-value of differential expression. **Supplementary table** 1 provides more details regarding the replicate group for each sample, and treatment and control information for each differential contrast. Next, we generated a feature matrix for each differential contrast. In our analysis, feature matrix was constructed by two steps. First, for each differential contrast, we generated a list of DEGs as positive training examples and sampled equal number of negative examples from the genome. The feature matrix *X* is a *n* by *m* matrix where *n* is the sum of number of positive examples and negative examples, and *m* is the number of TFS. In the second step, information from expression data, DAP-seq data and ATAC-seq data were integrated to construct *X.* Each entry *X*_*ij*_ in the feature matrix is computed by the following equation:

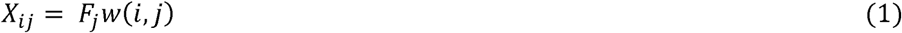

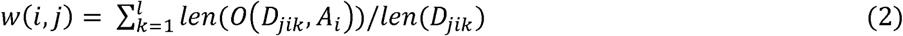

Where *j* denotes *j*th TF and *i* denotes *i*th gene (either positive or negative gene). *F*_*j*_ is the log2 fold change value of TF *j*. In equation (1), *w*(*i,j*) is the weight for each *X*_*ij*_. In equation (2), *D*_*jik*_ denotes the *k*th DAP-seq peak region of TE *j* found in the promoter region of gene *i*. The weight *w*(*i,j*) was computed by summing all *l* DAP-seq peak regions of TF *j* found in the promoter region of gene *i*. We evaluated each DAP-seq peak region by information from ATAC seq, which was done by searching overlapping regions between each DAP-seq peak on gene *i* and all open chromatin regions on gene *i* (denoted by *A*_*i*_), and the sum of length of overlapping regions was divided by the length of DAP-seq peak. This integration method will give higher weight *w*(*i,j*) if DAP-seq peaks for a TF j have more overlapping regions with the open chromatin regions found on gene i. *w*(*i,j*) equals zero if no DAP-seq peaks of TF *j* can be found on gene *i* or no ATAC-seq peaks can be found on gene *i* or no overlapping regions were detected between them.

To efficiently search for all overlaps, we constructed an interval tree for ATAC-seq peaks in each chromosome then iterated over each DAP-seq peak to find all overlaps between DAP-seq peak and ATAC-seq peaks. Python package Intervaltree (https://github.com/chaimleib/intervaltree) was used to perform the search. While our current analysis only explored the use of DAP-seq interaction data and ATAC-seq open chromatin region data, other types of interaction data and chromatin feature data can be easily integrated into equation 1 and equation 2. We will leave this to future exploration.

To construct feature matrices with only DAP-seq data, we marked each entry *X*_*ij*_ by ‘1’if binding site(s) of TF *j* are found in promoter region of gene *i* and ‘0’ if not.

We normalized the feature matrices by min-max normalization. Each *X*_*ij*_ was normalized by:

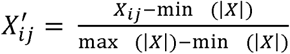

Where min (|*X*|) is the smallest absolute value in feature matrix *X* and max (|*X*|) is the largest absolute value in feature matrix *X*. During cross-validation, we computed min (|*X*|) and max (|*X*|) from training feature matrix and used them to normalize validation feature matrix and testing feature matrix.

## Machine learning models and feature selection

We tested several machine learning methods for classification, including logistic regression (LR), support vector machine (SVM), random forest (RF) and deep neural network (DNN). To perform feature selection, we applied different regularization techniques to each classifier. The details of classification and feature selection methods are described below.

### LRLASSO

This method is logistic regression with lasso penalty, which uses L1 regularization for feature selection (Lee et al. 2006). LRLASSO minimizes the following loss function:

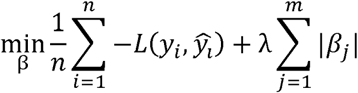

Where *y*_i_ and ŷ_*l*_ are the true label and predicted label for each training example, respectively. ŷ_*l*_ is estimated by the logistic function:

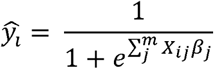

L(*y*_i_,ŷ_*l*_) is the log likelihood function and 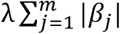 is the L1 penalty term.*β*_*j*_ is the coefficient for feature *j* (In our analysis, TF *j*). L(*y*_i_,ŷ_*l*_) is usually calculated by the cross-entropy loss function:

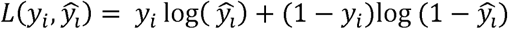

To perform feature selection, we tuned the L1 penalty parameter λ for this model using R package gglasso (Yang and Zou 2015). Given a sequence of ordered λ values, gglasso computes the solution for each λ iteratively. The computed solution for current λ will be used as initial value for next λ in the sequence. For each round of cross-validation, we used a sequence of 100 λ values which ranged from min(λ) to max(λ) and were spaced evenly on a log scale. max(λ) is the smallest λ value that shrinks all coefficients to zero. And min(λ) = η * max(λ), where η is a factor specified by user. For more details, please see the publication of gglasso (Yang and Zou 2015) and the online documentation of the R pacakge (https://cran.r-project.org/web/packages/gglasso/gglasso.pdf). In this way, each λ generates a LRLASSO model by training on training data set and the model was evaluated using validation data set to determine which λ gave the best prediction accuracy. Then the λ and the model with best prediction accuracy was again evaluated by the test data set.

### LGLASSO

This method is logistic regression with group lasso penalty. The group lasso penalty regularizes the coefficients, *β*_1_, *β*_2_…*β*_*m*_ by grouping and summing them using L2 norm (Meier et al. 2008; Yang and Zou 2015). If the *m* features can be grouped into *l* groups, the loss function of LGLASSO can therefore be written as:

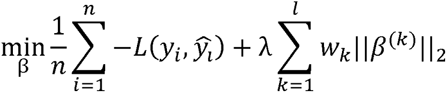

Where ||*β*^(*k*)^||_2_ is the L2 norm of all *β* s in group *k* and *w*_*k*_ is the weight for group *k*. We followed the default choice in gglasso paper (Yang and Zou 2015). We set 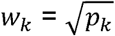, where, *p*_*k*_ is the number of features in group *k*. To obtain the prior grouping information for TFs, we ran CoReg (Song et al. 2017) on DAP-seq interaction network to identify co-regulator groups and used these groups in LGLASSO. For hyperparameter tuning and search, we used the same approach with LRLASSO.

### LREN

This method is logistic regression with elastic net penalty, which uses L1 + L2 regularization for feature selection. The loss function for LREN can be written as:

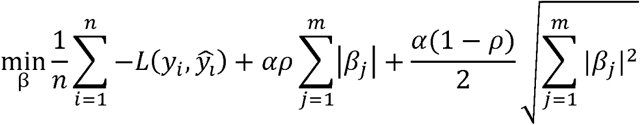

Where 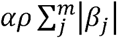 is the L1 penalty term and 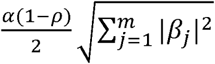 is the L2 penalty term. The parameter *α* and *ρ* control the strength of L1 penalty and the ratio of L1 penalty, respectively. Therefore, these two parameters are the only hyperparameters for LREN. For *α*, we used a sequence of five *α* values which range from 10^−3^ to 10^3^ and are evenly spaced on a log scale. For *ρ*, we used 0, 0.25, 0.5, 0.75 and 1 for tuning. Then grid search was performed to find the best combination of *α* and *ρ*. We used the function SGDClassifier() from scikit-learn package to perform training for LREN.

### LRPCC

This method is logistic regression with Pearson correlation coefficient (PCC). We computed the PCC between each feature and the true labels across all training examples. These features were then ranked using the PCC values by descending order. Similar to the process of stepwise linear regression, we iteratively added top *k* features at a time to train the model (Draper and Smith 1981) and the best set of features were determined by calculating accuracy of the trained model on validation data set. We set *k* as 50 to efficiently train the model. We used the function LogisticRegression() from scikit-learn package to perform training for LRPCC.

### GRRF

This method is guided regularized random forest. GRRF first trains a RF model and then uses the importance scores computed from this model to guide the feature selection process (Deng and Runger 2013). Briefly, GRRF computes a normalized importance score for each feature based on the original importance score from RF model. GRRF also assigns a penalty coefficient *λ*_*i*_(*λ*_*i*_ ∈ (0,1]) to each newly introduced feature when Gini information gain is calculated to split tree nodes. This is to penalize the use of new feature if it has not been used for splitting the node. *λ*_*i*_ is calculated by:

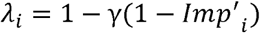

Where γ is the only hyperparameter that controls the strength of regularization. For more details about the algorithm of GRRF, please see the reference (Deng and Runger 2013). Since GRRF is more computationally expensive than other linear model-based methods and SVM-based method, we only explored γ = 0, 0.5 and 1 for hyperparameter tuning.

### LSVM

This method is linear support vector machine with L1 regularization. The only hyperparameter for this model is C, which controls the strength of L1 regularization. Smaller C will apply stronger regularization to the model, leading to a sparser model with many coefficients shrunk to zero. We used the function LinearSVC() from scikit-learn to perform training for LSVM. In our analysis, we used a sequence of C values which ranges from min(C) to 10^3^ and are evenly spaced on a log scale. min(C) is the minimum value of C which ensures all coefficients will be non-zero. min(C) is computed by calling the function l1_min_c() from scikit-learn library. Then the best C was determined by training the model on training data set and evaluating on validation data set.

### DNN

This method is deep neural network with L1 regularization for feature selection. The use of regularized DNN for genomic feature selection has been investigated in a previous publication (Li et al. 2015). The authors added a one-to-one layer between the input layer and hidden layers. L1 and L2 regularization were applied to the one-to-one layer to select features. Due to high computational cost of tuning hyperparameters of DNN, we chose to use only L1 regularization in the one-to-one layer and hidden layers. We used a similar DNN architecture with the previous publication (Li et al. 2015). In the input layer, there are 387 neurons and this number is equal to the number of input features (TFs). In the second layer (one-to-one layer), the same number of neurons are used and each is connected to one neuron from the input layer. Then we added two hidden layers which have 32 and 16 neurons after the one-to-one layer. The first hidden layer is fully connected with one-to-one layer and second hidden layer is fully connected with the first hidden layer. The last layer is an output layer which only has one neuron. Batch normalization was applied to one-to-one layer and each hidden layer to accelerate the training process. See (Li et al. 2015) for more details about using DNN to select features.

For hyperparameter tuning, we tuned L1 regularization parameter λ for DNN model. We used a sequence of 10 λ values which range from 10^−6^ to 10^3^ and are evenly spaced on a log scale. Adam optimizer was used to train the DNN model and learning rate α was fixed as 0.1. We compiled and trained DNN model using Keras library (https://keras.io/) with CUDA GPU acceleration. Training, hyperparameter tuning and testing was performed in the same way as described in other methods.

We provide ConSReg as a Python library for model training, tuning and testing using the models described in this study (github link https://github.com/LiLabAtVT/ConSReg).

## Evaluation strategy

### Evaluating different conditions

To evaluate the effect of selecting negative training examples, we tested three different methods: 1) non-significantly differentially expressed genes (NDEGs), which have p-value > 0.05 2) low-expressed genes (LEGs), which have mean expression between 0 and 0.5. 3) undetected genes (UDGs), which have mean expression value equal to zero. To evaluate the effect of promoter region length, we constructed feature matrices using three different promoter lengths, which are 1) 5kb upstream of TSS to 1kb downstream of TSS; 2) 3kb upstream of TSS to 0.5kb downstream of TSS; and 3) 0.5kb upstream of TSS to TSS. The promoter region length is passed as input argument to ChIPseeker package to search for corresponding genes for each DAP-seq peak. To evaluate the effect of methylated DAP-seq peaks VS all available DAP-seq peaks, we constructed the feature matrices using the DAP-seq peaks from regular DAP-seq (methylated DAP-seq peaks) and the DAP-seq peaks from amp-DAP-seq, where methylation was removed. The performance of two methods were then compared.

### Cross-validation for the models

For each feature matrix, we split the matrix into three subsets: 60% for training, 20% for validation (hyperparameter tuning) and 20% for testing. We trained the machine learning models on training data set and found the optimal set of hyperparameters by evaluating the trained model on validation data set (**Figure 1B**). Then the final performance of model with optimal hyperparameters was evaluated using the test data set. We used AUC-ROC and AUC-PRC as the metrics for evaluation. This process was repeated five times for each feature matrix to obtain the mean and standard deviation of AUC-ROC and AUC-PRC.

### Compare performance to enrichment-based method

We compared our methods to enrichment-based method. Similar to the approach used in TF2Network (Kulkarni et al. 2017), we computed the statistical significance of enrichment for each individual TF by hypergeometric test. The probability mass function is defined as:

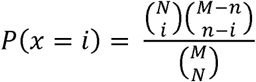

Where each parameter is explained below:

*i* is the number of DEGs that have DAP-seq peak(s) of the current TF.

*N* is the total number of DEGs in the current differential contrast.

*n* is the total number of protein-coding genes that have DAP-seq peak(s) of the current TF.

*M* is the total number of protein-coding genes.

p-values for all TFs were then computed by hypergeometric test and corrected by Benjamini-Hochberg correction (Y.Hochberg 1995). We used the same training set of positive genes and negative genes to compare LRLASSO with enrichment-based method. For each condition, this training set is the same feature matrix we used to evaluate machine learning models

To calculate AUC-ROC value for the enrichment-based method, we first ranked all TFs by ascending order using corrected p-values. Then we iterated over the ranked list of TFs. In each iteration, we used top *k*TFs as predictors and *k* is increased by one in next iteration until all 387 available TFs were included as predictors. Gene is considered as predicted positive if it has any predictors’ peak regions in its promoter region and predicted negative if not. Therefore, false positive genes are those predicted as positive but are actually negative in the training set and false negative genes are those predicted as negative but are actually positive in the training set. We calculated false positive rate 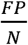 and false negative rate 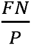 in each iteration and then all points of 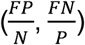 were put together to form the ROC curve for computing AUC-ROC value.

Since hold-out test will not be applicable for enrichment-based method, for both LRLASSO and enrichment-based method, training and testing were performed using the same training set to have fair comparison.

## Ranking TFs by stability selection

Since coefficients generated by LRLASSO model do not reflect the importance of each TF and the selected set of TFs would be slightly different when coefficients are initialized randomly. We applied stability selection (Meinshausen and Bühlmann 2010) to generate robust feature selection result from LRLASSO.

Randomized lasso was proposed as an implementation of stability selection for lasso method. The difference between randomized lasso and regular lasso is that subsampling of training examples and random perturbations for features are introduced into the feature selection process (Meinshausen and Bühlmann 2010). Briefly, a subset of training examples were selected and their features were randomly perturbed. Then a lasso model was trained using the perturbed subset of original training data set. This process was then repeated multiple times. The idea is that important features will be selected more often than the unimportant ones during this randomized process. When used with LRLASSO, the objective function of randomized lasso can be written as:

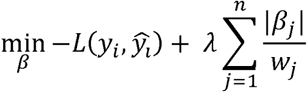

Where L(*y*_*i*_, ŷ_*l*_) is the log likelihood function as described previously in **Methods** section. *β*_*j*_ is the coefficient for feature *j*. 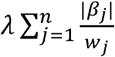 can be considered as the penalty term for randomized lasso, similar to L1 penalty term for regular lasso model. The only difference here is that random perturbation is introduced by *w*_*j*_, a scaling factor sampled from the range (0,1]. For simplicity of implementation, features can be rescaled to have the same effect with rescaling the coefficients (Meinshausen and Bühlmann 2010).

In our analysis, we randomly sampled half of the training examples from a feature matrix. Features were randomly perturbed by a scaling factor randomly sampled from (0,1]. Randomized lasso was performed *n* times for each feature matrix. For each feature, the final feature score was calculated as number of times the feature gets non-zero coefficient divided by *n*. In our analysis, we set *n* = 200.

## Network inference

GRN was inferred by computing the score for each TF using LRLASSO + stability selection and connecting the selected TFs to corresponding target genes. The detailed steps are described in this section.

First, we computed importance score for each TF in each differential contrast. There could be multiple feature matrices available for a single condition and each feature matrix corresponds to a differential contrast between treatment group and control group. For each of these differential contrasts, LRLASSO + stability selection was run and TFs with importance score > 0.5 were selected as regulators in GRN.

Second, with the selected regulators for each differential contrast, we constructed the GRN. If the selected TF j has a non-zero entry *X*_*ij*_ in feature matrix *X*, TF j is considered as having impact on target gene i. Then TF j will be linked to target gene i. To capture the most active regulatory interactions, we limit target genes to be only DEGs in each differential contrast. This process would generate multiple networks and each of them corresponds to a differential contrast. We then merged all networks generated in a single condition to build a final GRN for that condition.

